# U.S. imperiled species are most vulnerable to habitat loss on private lands

**DOI:** 10.1101/556241

**Authors:** Adam J. Eichenwald, Michael J. Evans, Jacob W. Malcom

## Abstract

To stem the ongoing loss of biodiversity, there is an urgent need to distinguish between effective and ineffective approaches for protecting species habitats. Using Google Earth Engine and 30 years of Landsat images, we quantify habitat change for 24 vertebrates on the U.S. Endangered Species List (ESA) and IUCN Red List across different categories of land ownership (e.g., federal, state, and private) that are subject to different conservation-focused legal restrictions. These estimates exclude changes due to agricultural conversion and burned areas. Imperiled species evaluated lost the least habitat (3.6%) on federal lands, while losses on private lands without conservation easements were more than twice as great (8.1%). Differences in annual percent loss before and after species were placed on the Endangered Species List, and between ESA and Red List species suggest that the ESA limited habitat loss and was most effective on federal lands. These results emphasize the importance of federal lands for protecting habitat for imperiled species and highlight the need to improve habitat protection on private lands for long-term conservation.

---

Habitat destruction is the primary driver of global biodiversity loss, and reducing or reversing habitat loss is a critical goal of conservation (Betts *et al.* 2017). This is particularly vital for the conservation of imperiled species, because habitat loss negatively affects species’ population size (Donovan and Flather 2002) and reproductive success (Kurki *et al.* 2000). Given the current global biodiversity crisis (Betts *et al.* 2017), it is critical to identify drivers and effective mechanisms for preventing future loss. Therefore, rigorous analyses that evaluate the effectiveness of conservation laws are needed to identify successes and shortcomings in efforts to maximize habitat protection and inform policy decisions.

The United States has some of the strongest laws in the world protecting imperiled species but can regulate federal and non-federal entities differently. The Endangered Species Act (ESA) provides a prominent example. The ESA prohibits the take (e.g., harm, kill, harass) of protected animals but implements this protection differently for federal and non-federal actors. Incidental take of protected animals as a result of federal actions is authorized under Section 7, which is widely used (Malcom and Li 2015; Evans et al. 2019). In contrast, non-federal entities receive authorization for incidental take of listed animals through section 10, a mechanism that is difficult to enforce (Carter 1991). While both processes are initiated by the regulated party, Section 7 consultation is standard practice for federal agencies and federal decisions that trigger consultation are publicly visible; by contrast, it is unknown how often private entities eschew the Section 10 process. Furthermore, federal land management agencies have specific legal and regulatory conservation mandates (e.g., Federal Land Planning and Management Act, the National Forest Management Act) that protect imperiled species. Thus, protections for imperiled species may be stronger on federally owned lands.

The question remains as to whether differences in conservation law implementation between federal and non-federal contexts result in observable differences in protections for imperiled species habitat. Weaker protections on non-federal land would severely limit conservation because so much biodiversity exists outside of protected areas (Jenkins *et al*. 2015). However, scientists have lacked the data and tools to evaluate patterns of imperiled species habitat loss at large scales. Instead, research has focused on small-scale case studies (Smith *et al.* 2018). Such studies of limited geographic or taxonomic scope are difficult to extrapolate beyond their study sites. While informative, these results are insufficient for informing policy debates at the national scale on effective approaches to biodiversity conservation. Large-scale data and analyses are needed to inform policy and enable oversight of land management agencies (Trainor *et al.* 2013).

Advances in remote sensing and cloud computing now enable the rapid analysis of global satellite imagery datasets, allowing large-scale, long-term analysis of habitat loss (Song *et al.* 2018). Here we use satellite imagery to report on 30 years of habitat loss for 16 ESA listed species and 8 listed species on the International Union for Conservation of Nature’s Red List (IUCN 2018) in different land ownership classes in the contiguous US, not including agricultural land. Unlike ESA listed species, species on the Red List are not legally protected. Differences in protections among species and between federal and non-federal lands creates an opportunity to evaluate how effectively conservation policies in the United States protect imperiled species habitat, and to identify strengths and weaknesses in their implementation. Our goal was to quantify differences in habitat protection for imperiled species between federal and non-federal lands and to evaluate the degree to which ESA contributes to habitat protection.

## Methods

### Study Species and Area

We measured habitat loss for 24 vertebrates in the contiguous US from among those listed under the ESA and on the Red List (Figure 1).We selected species with ranges modeled by the USGS Gap Analysis Program (GAP) that contained both federal and non-federally owned land. GAP models provide a methodologically consistent representation of species ranges at 30m resolution, which are built using a national wildlife habitat relational database based on habitat associations described in published literature (including detailed land cover, elevation, and hydrological characteristics). We used these models to delineate the areas that represented where each species was most likely to be found. The GAP program provides maps of suitable habitat for over 2000 listed and unlisted species in the US, most of which have limited ranges that were less useful for our goals (USGS 2018). The 24 species that we selected were those with ranges that encompassed both federal and non-federally owned land, that relied on habitats that could be detected in satellite imagery, and collectively inhabited all major ecoregions in the continental US.

**Figure 1.**
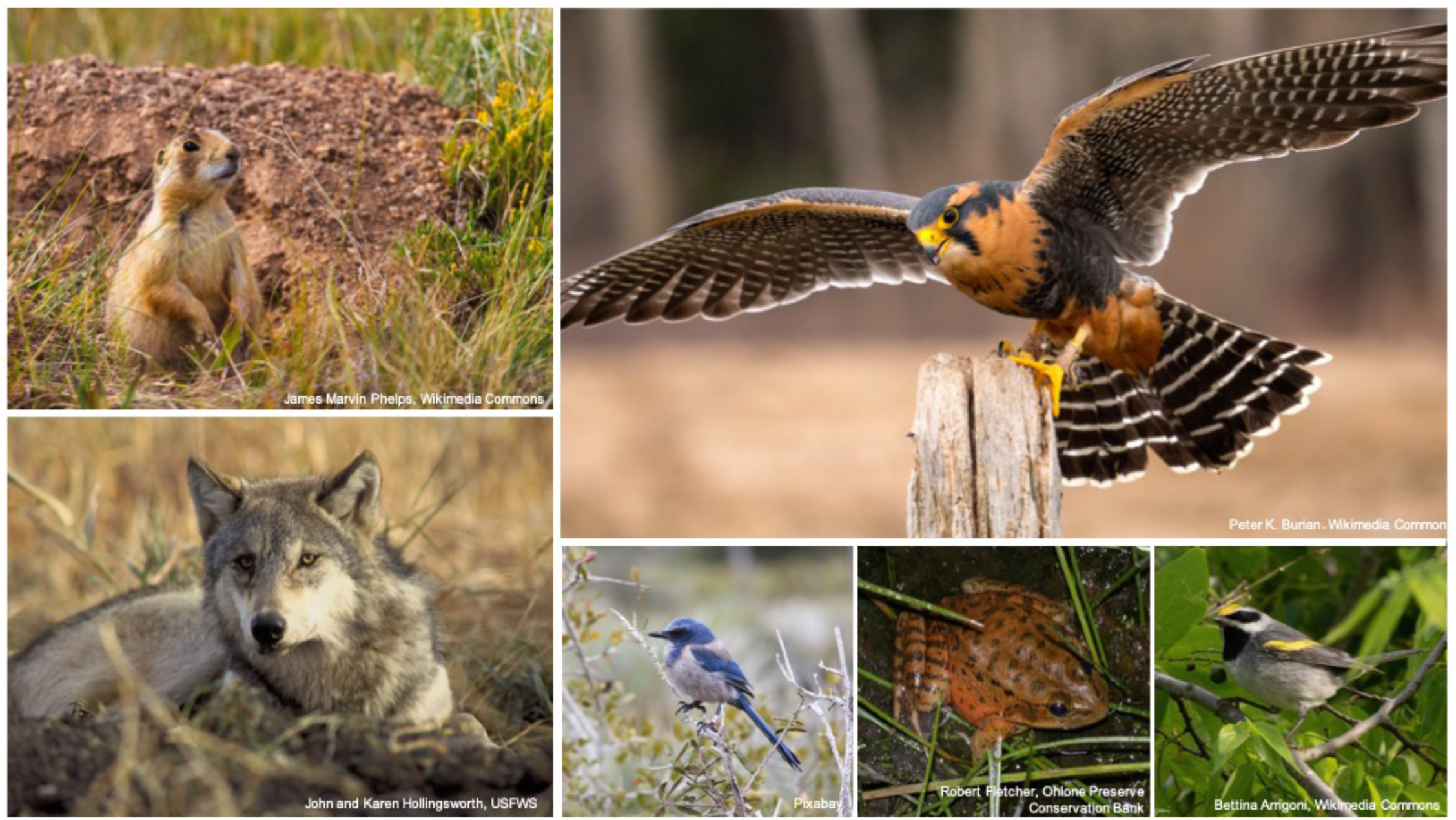
Some of the listed species used in our analysis. Clockwise from the top left: Utah prairie dog (*Cynomys parvidens*), aplomado falcon (*Falco femoralis*), golden winged warbler (*Vermivora chrysoptera*), California red legged frog (*Rana draytonii*), Florida scrub jay (*Aphelocoma coerulescens*), and gray wolf (*Canis lupus*).

For ESA listed species, we also considered an additional delineation of species range: the legally-recognized ranges documented and provided by FWS (“administrative range,” https://ecos.fws.gov). These administrative ranges vary in how they are designed. Many only represent presence/absence of the species at county level resolution, while others may be delineated using other readily available geological/ecological features like watersheds. As GAP ranges are not legally recognized, we include both range representations so that we can compare habitat loss within all areas that species may be ecologically likely to occur (i.e., GAP range) to the subset of areas receiving federal protections (i.e., administrative range). This comparison helps further evaluate the impact of federal protections on habitat loss by defining the explicit species ranges where the ESA is legally applicable.

To delineate land ownership classes, we used the US Protected Areas Database v 2.0 (PAD, http://gapanalysis.usgs.gov/padus) and classified lands as federal, state, NGO, protected private, or non-protected private. We intentionally did not include tribal lands in our analyses, as they are legally governed by separate nations. In aggregate, these species represented multiple ecoregions, covering 49% of the contiguous US (WebTable 1).

### Remote Sensing Analysis

We used the Google Earth Engine implementation of the LandTrendr algorithm (Kennedy *et al.* 2018) to identify loss of imperiled species habitat between 1986 and 2018 from Landsat imagery via breakpoints in temporal trends of NDVI. We defined habitat loss as the area where one habitat was degraded quickly over a short period of time. We did not consider landscape metrics like fragmentation or connectivity in our analyses. We standardized thresholds used to identify breakpoints among habitat types using the mean and standard deviation within each focal species range (Yang *et al.* 2018). We discarded disturbances less than 450m^2^ to reduce oversensitivity. Agricultural crop rotations can confound change detection by creating false positives. Therefore, we masked LandTrendr output using the USDA 30m resolution Cropland Data Layer from 2017 (Boryan *et al.* 2011), excluding agriculture conversions prior to 2017. This created a conservative bias to loss estimates.

LandTrendr is also likely to detect habitat loss from fire. However, because we could not distinguish between natural and prescribed burns, fire would confound our assessment of legal and regulatory protections. Therefore, including burned areas up to the year they were burned in our analysis may have introduced a confounding variable. We eliminated burned areas using data from the Monitoring Trends in Burn Severity Project (MTBS Project 2018). The percent land excluded due to fire and agriculture per species is shown in Webtable 2.

### Habitat Loss Analysis

We calculated annual percent habitat loss as the proportion of pixels within a species’ range in each land ownership category showing habitat loss in each year. Total pixels within the range remained constant across years. To account for background trends in habitat loss, we standardized annual percent loss estimates by subtracting the mean annual percent loss across all species and land ownership types in a given year from species-specific annual percent loss estimates. We refer to this measure as “adjusted annual loss.” We also calculated total percent loss as the proportion of pixels over the entire period that were disturbed and did not recover back to their original NDVI values (total percent loss). We used the LandTrendr algorithm to identify breakpoints resulting in NDVI gain rather than loss; total percent loss only includes pixels that were disturbed and never regained NDVI in subsequent years. We excluded years corresponding to the beginning of our data collection (1986) and transitions between Landsat satellites (2001), because these years exhibited extreme peaks.

To identify important predictors of habitat loss, we fit linear mixed models estimating adjusted annual loss per species as a function of land ownership (“Zone”), time (“Year”), listing status (“Status”), and whether the GAP or FWS range is considered (“Range”). All models included a random intercept per species to account for correlation in repeated measures. Unless otherwise specified, we consider losses for ESA listed species within the FWS range that occurred when the species was listed. Models were estimated in a Bayesian framework using the rstanarm package (Goodrich *et al.* 2018) for R (RCore Development Team 2012). We used default priors and sampled 1,000 iterations of four Markov chains following a 1,000 iteration burn-in period. Chain convergence for all parameters was assessed using the 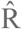 statistic, with 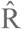 < 1.1 indicating convergence (Gelman and Rubin 1992). We used the Watanabe-Akaike Information Criterion (WAIC) to measure support among competing models (Watanabe 2010). We used the sum of WAIC weights (ω_i_) among models including a given variable to measure variable importance, and use Σω_i_ > 0.90 to signify important variation in loss as a function of a given variable (Watanabe 2013).

To test for differences in habitat loss among land ownership classes, we fit a candidate set of linear mixed models that included an intercept-only (null) model and all combinations of the effects of Year, Zone, and Status. This candidate set was fit to adjusted annual loss data from ESA-listed and IUCN species within FWS and GAP ranges, respectively. To test for differences in habitat loss inside and outside administrative ranges, we also fit an identical model set with Range substituted for Status to adjusted annual loss data for ESA-listed species from within GAP distribution ranges inside and outside of the FWS range, substituting Range for Status. Because Red List species do not have administrative ranges, the candidate sets were fit to different data and evaluated separately. Additionally, we tested for differences in percent loss among pairs of land ownership types, between ESA listed and Red List species, and between ranges using the maximum probability of effects (*MPE*) and 95% credible intervals around pairwise contrasts.

As a post hoc analysis, we also estimated differences in habitat loss trends over time among land ownership classes. We considered four trends in adjusted annual loss representing four simple, potential patterns of change: linear, logarithmic, exponential and quadratic. To identify the appropriate trend form, we fit a set of four mixed models to adjusted annual loss data as a function of year separately for each by ownership class to account for potential differences in the form of trend among classes. We identified the most supported relationship as the model receiving the lowest WAIC score. We compared trends among land ownership types in terms of the form of the trend that was most supported, and whether slopes were positive, negative, or effectively zero. This analysis was conducted for ESA-listed and Red List species separately.

Finally, we evaluated three predictions to test the hypothesis that the ESA was the mechanism responsible for observed differences in habitat loss. If the data are consistent with the hypothesis, we expect:

1. Lower adjusted annual loss within ESA listed species ranges after they were listed relative to when they were listed.
2. No differences in total percent loss among land owned by different federal agencies. Differences would indicate agency-specific regulations as the mechanism driving loss reduction.
3. Lower total percent loss for ESA listed species than for Red List species.

We tested expectation #1 by estimating the effect of ESA listing on adjusted annual loss using a linear mixed model with a random intercept per species and fixed effects on the interaction between an ESA listing indicator variable and Zone. For expectation #2, we used a linear mixed model estimating differences in total percent loss within the FWS range of listed species on federal lands as a function of the federal agency that owned the land. For expectation #3, we used the effect of listing status on total percent loss in GAP ranges for Red List species and in FWS ranges for ESA listed species. For ESA listed species, we considered only losses after listing. In all analyses, we inferred differences among groups and meaningful trends if the relevant coefficient had an *MPE* > 95% and a 95% credible interval around estimated effect sizes that did not include zero.

## Results

Variation in adjusted annual habitat loss for imperiled species was explained by land ownership types, listing status (ESA vs.Red List), the range considered (ecological vs. administrative), and across years. Linear mixed effects models containing a three-way interaction between Zone, Time, and Status or Range received the most support (ΔWAIC > 13.9; WebTable 3). Imperiled species lost the least habitat on federal lands (u = 3.6%, sd = 3.8), significantly less than all other types (Δ ≥ 6.2%, MPE ≥ 0.97). Species lost the greatest amount of habitat on non-protected lands (µ = 8.6%, sd = 6.3). Total loss on both non-protected and protected private lands was significantly greater than all other ownership types (Δ ≥ 15.0%, MPE = 1.00; Figure 2). There was no difference (Δ ≤ 1.5%, MPE = 0.69) between net loss on NGO (µ = 4.5%, sd = 3.8) and State (µ = 4.6%, sd = 3.6) lands. Additionally, total loss was higher within ecological ranges than administrative ranges (Δ ≥ 14.4%, MPE = 1.00; Figure 2).

**Figure 2.**
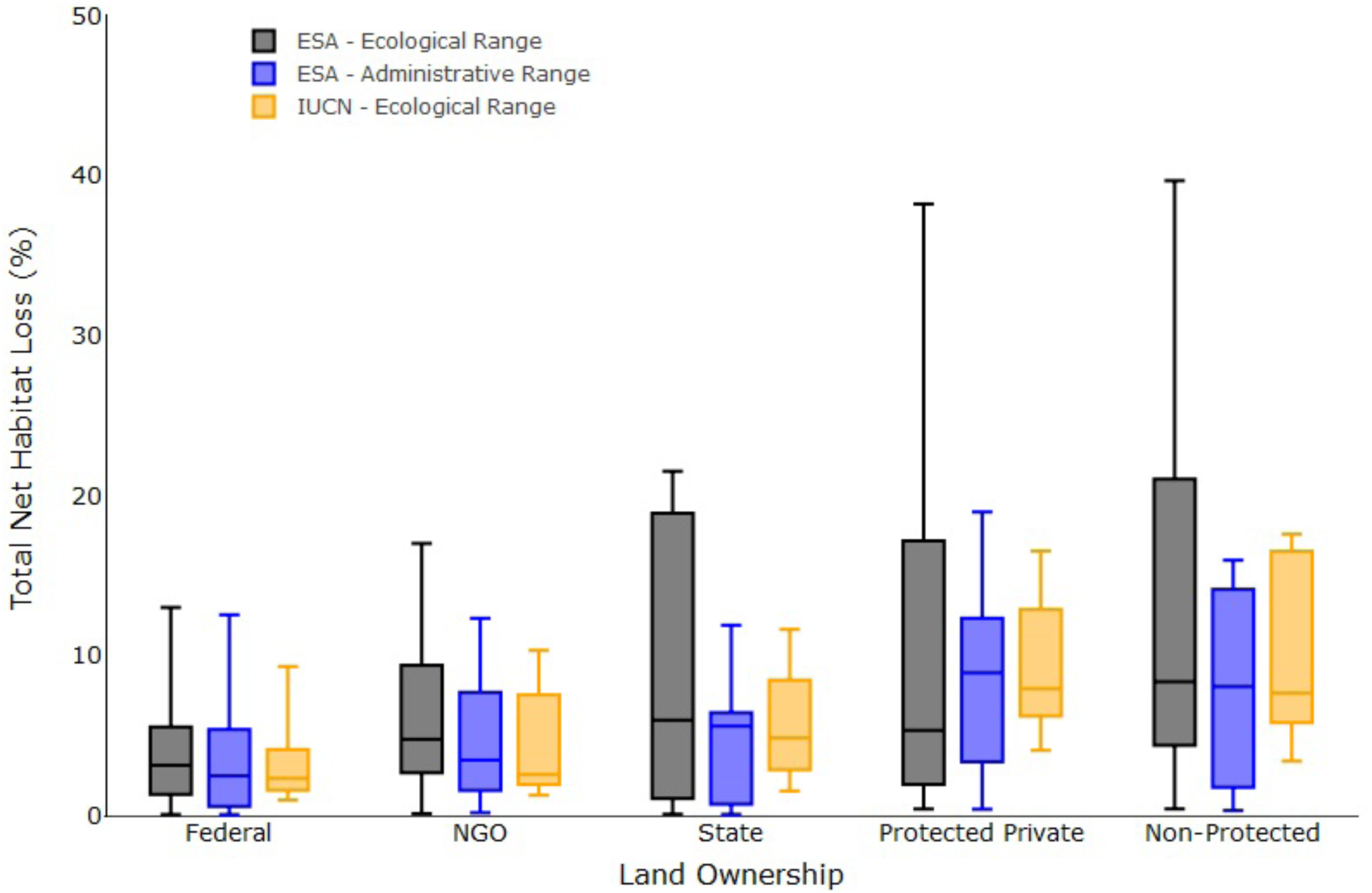
Rates of habitat loss within imperiled species ranges were lowest on Federal lands, and highest on Non-Protected private lands. Box plots show the distribution of net habitat loss among imperiled species from 1986 to 2017. Percentages differed for ESA listed species (ESA) depending on whether the potential/ecological range (black) or administrative range (blue) was considered. Total percent losses within Red List species ranges (orange) were not significantly greater than those for ESA listed species according to our prespecified Bayesian criteria, but they did lose marginally more habitat overall.

Trends in annual adjusted loss over time also differed among land ownership types (Figure 3). A linear decline was the best supported model of annual habitat loss on NGO lands for both listed and Red List species, indicating consistently decreasing loss rates (WebTable 4). Annual habitat loss increased logarithmically on both protected and non-protected private lands for listed and Red List species, indicating increasing annual loss that stabilized over time (Figure 3). Among Red List species, there was no change in annual loss over time on state or protected private lands (WebTable 4). Annual loss of habitat for listed species on state lands increased exponentially (WebTable 4). On federal lands, quadratic trends indicate decreasing annual loss rates from 1986 to 2005 for listed species, and 1986 to 2008 for Red List species, after which annual loss has been increasing (Figure 3).

**Figure 3.**
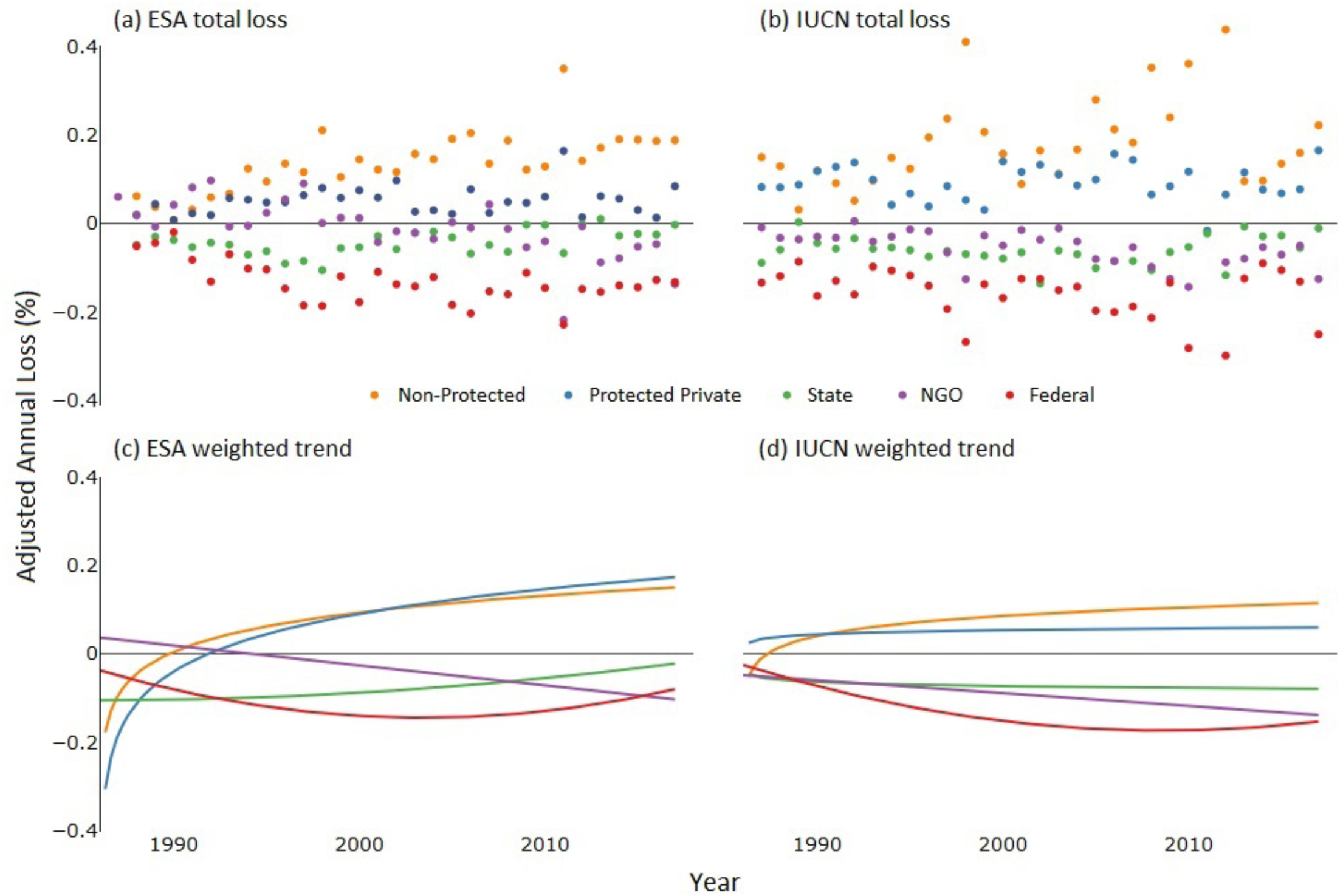
Federal and non-federal lands exhibited different trends in annual rates of habitat loss within imperiled species ranges. Data points (A, B) show the unweighted annual total percent habitat loss across species within each land management zone, adjusted by the mean loss in a given year. Trends over time were similar between (C) ESA listed species and (D) IUCN Red List species. Trend lines were estimated from the marginal relationship between year and adjusted annual loss accounting for random intercepts per species.

Species experienced significantly less habitat loss after they were listed compared to before (Δ = 25.0%, MPE = 1.00) (Figure 4). Total habitat loss was consistent among the six agencies managing federal lands in this study. Pairwise differences in total loss between agencies were not different from zero (MPE < 0.95), except that losses on National Oceanic and Atmospheric Administration lands (7,405,233.3 km^2^, all on beaches) were significantly lower than those on U.S. Bureau of Reclamation (2,896,025,400 km^2^) and Department of Defense (519,493.5 km^2^) lands (Δ ≥ 6.5%, MPE ≥ 0.96). ESA listed species lost marginally less habitat than Red List species overall (Δ = −2.62%, *MPE* = 0.67) and this difference was most pronounced on federal lands (Δ = −4.57%, *MPE* = 0.75).

**Figure 4.**
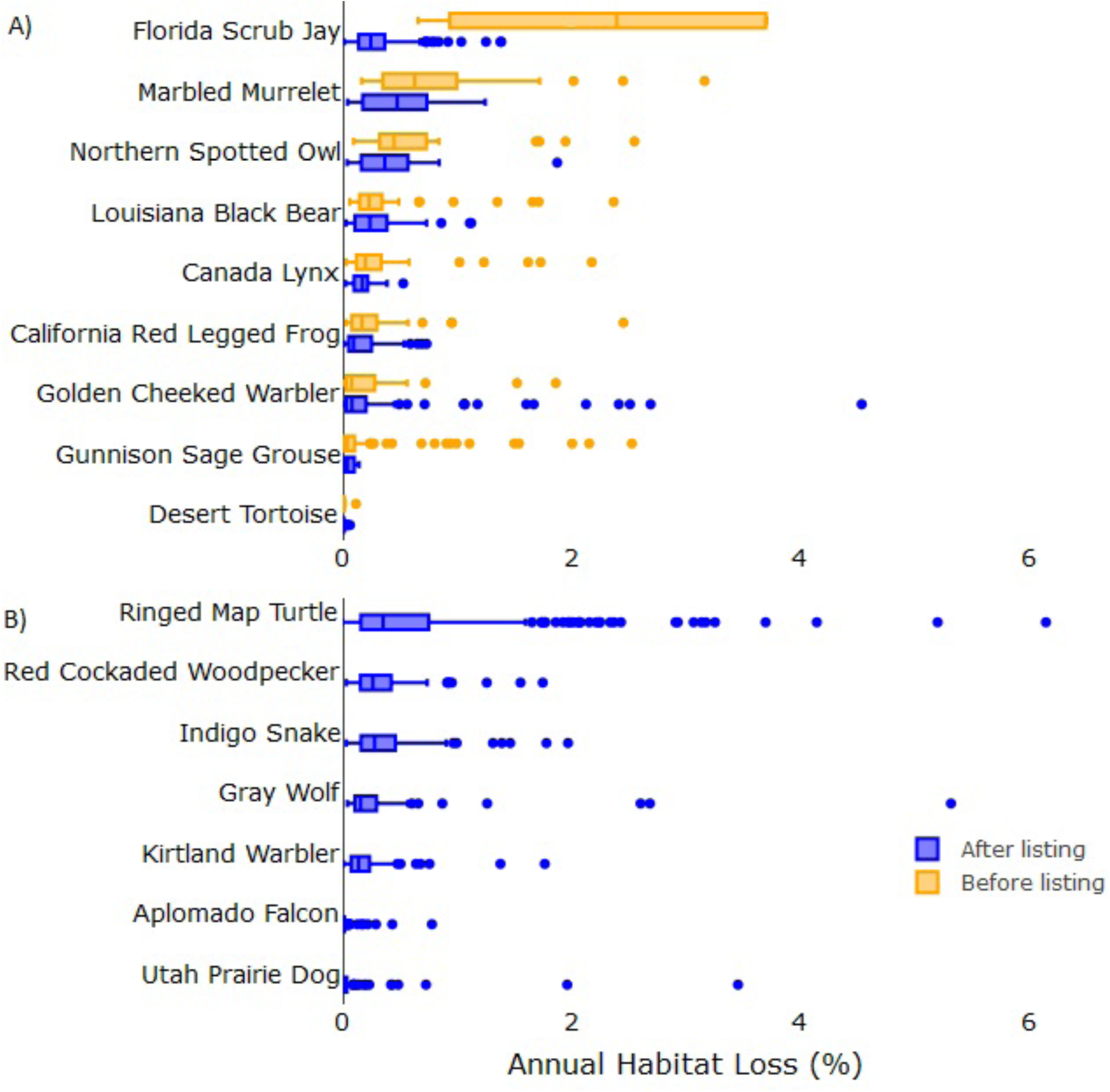
Species lost less habitat within their ranges while they were listed on the Endangered Species List. Box plots show the distribution of annual habitat loss for each ESA listed species as the percentage of species habitat within administrative ranges lost in each year before and after listing (A). Species listed prior to 1986 (B) did not have loss data before listing.

## Discussion

Critically evaluating approaches to conserving habitat is imperative to conserving imperiled species, yet historically scientists have only been capable of small-scale studies. We used satellite data to examine habitat loss in various land ownership classes across the United States to evaluate the efficacy of habitat protection in different land ownership contexts. Habitat loss for imperiled species was lowest on federal lands and highest on protected private and non-protected lands. Our results indicate that federal lands provide protections for imperiled species habitat and identify important shortcomings of protections outside of federally owned areas.

We found evidence that the ESA likely contributed to habitat protections on federal lands. Species lost less habitat after they were listed than before, and ESA species on federal lands also lost less habitat than Red List species. Consistent habitat loss among lands managed by different federal agencies indicates that agency-specific regulations (e.g., National Forest Management Act) were not primarily responsible for limiting loss. This further supports the hypothesis that legal protections under the ESA contributed to reduced habitat loss. The ESA is the only law in the United States that provides non-discretionary habitat protection across federal lands. While the National Environmental Policy Act requires federal agencies to report on the environmental impacts of planned actions, the law has no requirement to avoid, minimize, or mitigate these impacts. Therefore, while these laws almost certainly provide some degree of protection for imperiled species, our results are consistent with the hypotheses that the ESA provided unique protections across federal lands.

There are several reasons why the implementation and effectiveness of conservation laws are likely reduced on federal lands in comparison to private lands. First, individual property rights are highly protected in the U.S. (Knight 1999), and conservation laws like the ESA include exemptions for private actors (e.g., no protections for critical habitat on private lands, absent a federal nexus). Even where laws do apply, a lack of visibility and the voluntary nature of conservation initiatives on private lands mean that regulations may still fail to provide protections to habitat (e.g., oil development within Lesser prairie chicken range; Melstrom 2017). In addition to the lack of oversight over private actors, landowners may engage in preemptive habitat destruction to avoid perceived ESA land use restrictions (Lueck and Michael 2003; Brook *et al.* 2003), although such behavior is not guaranteed to occur (Melstrom 2017). Therefore, inefficient protections outside of federal lands will undermine present, past, and future conservation work.

Uneven protections are problematic to imperiled species conservation for three reasons. First, species do not only inhabit federally managed areas. In the U.S., there is a misalignment between biodiversity and federally protected areas (Jenkins *et al.* 2015), as more than half of listed species’ ranges include >80% private land (US Fish and Wildlife Service 2009). Not only did private lands show the highest rates of loss, our results likely also underestimate loss on private lands, as we do not account for loss from agricultural expansion. The cropland data layer showed that between 2008 and 2017, 127,100,181.6km^2^ (4.4%) of imperiled species habitat on private lands were converted to agriculture, while only 838,440km^2^ (0.1%) of federal lands were converted. This is consistent with national-level agricultural trends (Lark *et al.* 2015). Additionally, high and increasing losses on state lands indicate that current state protections are also inadequate. Relying on federal protections alone is therefore insufficient to conserve imperiled species.

Second, species within protected ranges can still be threatened by surrounding habitat loss. Isolated islands of protected habitat surrounded by development reduce the value of protected areas (Radeloff *et al.* 2010) and create extinction debts (delayed extinction following habitat loss) for species that might not be immediately noticed (Tilman *et al.* 1994). Furthermore, although habitats naturally change over time, ecosystems will transform dramatically with global climate change (Jones *et al.* 2009). Vertebrates are predicted to move their ranges in response, with a predicted 20% of US mammals shifting out of federal areas and into other land ownership classes (Burns *et al.* 2003). These interactions highlight the need for conservation laws to consider holistic landscape perspectives.

Finally, uneven protections likely limit species recovery. Protecting species only within administrative ranges potentially constrains species to current distributions instead of possible distributions (Carroll *et al.* 2010). Greater habitat loss within ecological ranges compared to administrative ranges are not surprising given that ESA interventions and other federal regulations are usually limited to these administrative areas, potentially reflecting an inability to protect unoccupied suitable habitat. Our results highlight that without extending protections to unoccupied habitat recovery may be hampered, especially for wide-ranging species.

This study examined a limited sample of the 1600 ESA listed species and does not include land covered by croplands or area burned by fire. Therefore, our results must be generalized with caution, particularly since our comparison of loss before and after ESA listing was limited to 8 species. Additionally, we selected species for which habitat loss is a significant threat, and our results cannot be used to infer the efficacy of legal protections against other threats. Furthermore, USGS GAP models were created in 2001. This may have resulted in the exclusion of habitat loss that occurred between 1986 and 2001. This exclusion presumably was unbiased by land ownership types and therefore should not change the results of our relative comparisons across land types. The exclusion of habitat loss prior to 2001 would affect analyses comparing habitat losses before and after listing for ESA species, leading to an underestimation of loss prior to listing. Therefore, the reduction in habitat loss for species after they were listed identified by our analysis may have been even greater than our results indicate (Figure 4).

It is important to note that our method is applicable to any species for which a discrete range can be identified, not just those modeled by GAP habitat distributions. We chose GAP models for consistency among species’ ranges, but researchers can apply the LandTrendr algorithm to measure habitat loss within any designated range for a given species.

Measurement and evaluation are critical for successful biodiversity protection. Our analyses indicate that federal protections have successfully reduced habitat loss for imperiled species in the U.S.. However, they also identify gaps that compromise wildlife conservation. To successfully recover imperiled species, the current differences in legal protections between federal and private lands must be minimized via increased enforcement, greater conservation incentives, or compliance assistance. All listed species experienced a net loss of habitat, even on federal lands, which may be symptomatic of systemic conservation challenges (Laurance 2010). Additionally, greater habitat loss among Red List species demonstrates that imperiled species without legal protection remain at risk. As species can become functionally extinct before losing 30% of their population (Saterberg *et al.* 2013), legally protecting species only once they face immediate threats may still result in extinctions (Rohlf 1991). As human well-being is intimately interconnected with biodiversity (IBPES 2019), a failure to protect imperiled species will have consequences beyond species extinctions.

## Acknowledgments

We thank J. Miller, M. Evansen, and M. Moskwik for assistance formulating project goals and analyses. J. Miller and J.M. Reed provided review and feedback that improved the manuscript.

---

Figure 1 Photo Attributions

Utah Prairie Dog: James Marvin Phelps. Flickr, https://www.flickr.com/photos/mandj98/28604441673

Aplomado Falcon: Peter K Burian, Wikimedia Commons, https://commons.wikimedia.org/wiki/File:Aplomado_Falcon_at_a_falconry_center.jpg

Florida Scrub Jay: https://pixabay.com/photos/florida-scrub-jay-bird-wildlife-1835209/

California Red Legged Frog: Robert Fletcher, Ohlone Preserve Conservation Bank, Flickr, https://www.flickr.com/photos/usfwsendsp/5840045028

Golden Winged Warbler: Bettina Arrigoni, Wikimedia Commons, https://commons.wikimedia.org/wiki/File:Golden-winged_Warbler_(male)_Sabine_Woods_TX_2018-04-28_06-59-11_(41300184395).jpg

Gray Wolf: USFWS Endangered Species, Wikimedia Commons, https://commons.wikimedia.org/wiki/File:Endangered_gray_wolf_(Canis_lupus).jpg

## Literature Cited

Betts MG, Wolf C, Ripple WJ, et al. 2017. Global forest loss disproportionately erodes biodiversity in intact landscapes. Nature 547: 441.

Boryan C, Yang Z, Mueller R, et al. 2011. Monitoring US agriculture: the US Department of Agriculture, National Agricultural Statistics Service, Cropland Data Layer Program. Geocarto International 26: 341–358.

Brook A, Zing M and De Young R. 2003. Landowners’ response to an Endangered Species Act listing and implications for encouraging conservation. Conservation Biology 17(6): 1638–1649.

Burns CE, Johnston KM and Schmitz OJ. 2003. Global climate change and mammalian species diversity in US national parks. Proceedings of the National Academy of Sciences 100: 11474–11477.

Carroll C, Vucetich JA, Nelson MP, et al. 2010. Geography and recovery under the US Endangered Species Act. Conservation Biology 24: 395–403.

Carter CH. 1991. A Dual Track Incidental Takings: Reexamining Sections 7 and 10 of the Endangered Species Act. Boston College Environmental Affairs Law Review 19: 73–134.

Donovan TM and Flather CH. 2002. Relationships among North American songbird trends, habitat fragmentation, and landscape occupancy. Ecological Applications 12: 364–374.

Evans MJ, Malcom JW and Li YW. 2019. Novel data show expert wildlife agencies are important to endangered species protection. Nature Communications 10: 1–9

Gelman A and Rubin DB. 1992. Inference from iterative simulation using multiple sequences. Statistical science 7: 457–472.

Goodrich B, Gabry J, Ali I, et al. 2018. rstanarm: Bayesian applied regression modelling via Stan. R Package Version 2.17. 4.

IBPES. 2019. IPBES Global Assessment. Intergovernmental Science-Policy Platform on Biodiversity and Ecosystem Services:

International Union for Conservation of Nature. 2018. The IUCN Red List of threatened Species.

Jenkins CN, Van Houtan KS, Pimm Sl and Sexton O. 2015. US protected lands mismatch biodiversity priorities. Proceedings of the National Academy of Sciences 112(16): 5081–5086

Kennedy RE, Yang Z, Gorelick N, et al. 2018. Implementation of the LandTrendr Algorithm on Google Earth Engine. Remote Sensing 10(5): 691.

Knight RL. 1999. Private lands: The neglected geography. Conservation Biology 13(2): 223–224

Kurki S, Nikula A, Helle P, et al. 2000. Landscape fragmentation and forest composition effects on grouse breeding success in boreal forests. Ecology 81: 1985–1997.

Lark, TJ, Salmon, JM and Gibbs, HK. 2015. Cropland expansion outpaces agricultural and biofuel policies in the United States. Environmental Research Letters, 10, 044003.

Laurance WF. Sodhi N and Ehrlich PR (Eds). 2010. Habitat destruction: death by a thousand cuts. In: Conservation biology for all. Oxford University Press.

Malcom JW and Li Y-W. 2015. Data contradict common perceptions about a controversial provision of the US Endangered Species Act. Proceedings of the National Academy of Sciences 112: 15844–15849.

MTBS Project. USDA Forest Service and U.S. Geological Survey (Eds). 2018. Fire Level Geospatial Data.

RCore Development Team. 2012. R: A language and environment for statistical computing. Vienna, Austria: R Foundation for Statistical Computing.

Saterberg T, Sellman S and Ebenman B. 2013. High frequency of functional extinctions in ecological networks. Nature 499: 468–470.

Smith JA, Brust K, Skelton J, et al. 2018. How effective is the Safe Harbor program for the conservation of Red-cockaded Woodpeckers? Condor 120: 223–233.

Song X-P, Hansen MC, Stehman SV, et al. 2018. Global land change from 1982 to 2016. Nature 560: 639.

Trainor A, Walters J, Urban D, et al. 2013. Evaluating the effectiveness of a Safe Harbor Program for connecting wildlife populations. Animal Conservation 16: 610–620.

US Fish and Wildlife Service. 2009. Our Endangered Species Program and how it works with landowners.

Watanabe S. 2010. Asymptotic equivalence of Bayes cross validation and widely applicable information criterion in singular learning theory. J Mach Learn Res 11: 3571–3594.

Watanabe S. 2013. A widely applicable Bayesian information criterion. J Mach Learn Res 14: 867–897.

U.S. Geological Survey. 2018. Gap Analysis Project Species Habitat Maps.

Yang Y, Erskine PD, Lechner AM, et al. 2018. Detecting the dynamics of vegetation disturbance and recovery in surface mining area via Landsat imagery and LandTrendr algorithm. Journal of Cleaner Production 178: 353–362.

